# Recurrent fusions in *PLAGL1* define a distinct subset of pediatric-type supratentorial ependymoma

**DOI:** 10.1101/2021.04.23.441059

**Authors:** Philipp Sievers, Sophie C. Henneken, Christina Blume, Martin Sill, Daniel Schrimpf, Damian Stichel, Konstantin Okonechnikov, David E. Reuss, Julia Benzel, Kendra Maaß, Marcel Kool, Dominik Sturm, Patricia Kohlhof-Meinecke, Ofelia Cruz, Mariona Suñol, Cinzia Lavarino, Viktoria Ruf, Henning B. Boldt, Mélanie Pagès, Celso Pouget, Leonille Schweizer, Mariëtte E. G. Kranendonk, Noreen Akhtar, Ulrich Schüller, Wolf C. Mueller, Hildegard Dohmen, Till Acker, Patrick N. Harter, Christian Mawrin, Rudi Beschorner, Sebastian Brandner, Matija Snuderl, Zied Abdullaev, Kenneth Aldape, Mark R. Gilbert, Terri S. Armstrong, David W. Ellison, David Capper, Koichi Ichimura, Guido Reifenberger, Richard G. Grundy, Nada Jabado, Lenka Krskova, Michal Zapotocky, Ales Vicha, Pascale Varlet, Pieter Wesseling, Stefan Rutkowski, Andrey Korshunov, Wolfgang Wick, Stefan M. Pfister, David T. W. Jones, Andreas von Deimling, Kristian W. Pajtler, Felix Sahm

## Abstract

Ependymomas encompass a heterogeneous group of central nervous system (CNS) neoplasms that occur along the entire neuroaxis. In recent years, extensive (epi-)genomic profiling efforts have identified several molecular groups of ependymoma that are characterized by distinct molecular alterations and/or patterns. Based on unsupervised visualization of a large cohort of genome-wide DNA methylation data, we identified a highly distinct group of pediatric-type tumors (n = 40) forming a cluster separate from all established CNS tumor types, of which a high proportion were histopathologically diagnosed as ependymoma. RNA sequencing revealed recurrent fusions involving the *pleomorphic adenoma gene-like 1* (*PLAGL1*) gene in 19 of 20 of the samples analyzed, with the most common fusion being *EWSR1:PLAGL1* (n = 13). Five tumors showed a *PLAGL1:FOXO1* fusion and one a *PLAGL1:EP300* fusion. High transcript levels of *PLAGL1* were noted in these tumors, with concurrent overexpression of the imprinted genes *H19* and *IGF2*, which are regulated by PLAGL1. Histopathological review of cases with sufficient material (n = 16) demonstrated a broad morphological spectrum of largely ependymoma-like tumors. Immunohistochemically, tumors were GFAP-positive and OLIG2- and SOX10-negative. In 3/16 of the cases, a dot-like positivity for EMA was detected. Consistent with other fusion-positive ependymal groups, all tumors in our series were located in the supratentorial compartment. Median age of the patients at the time of diagnosis was 6.2 years. Analysis of time to progression or recurrence revealed survival times comparable to those of patients with *ZFTA:RELA*-fused ependymoma. In summary, our findings suggest the existence of a novel group of supratentorial ependymomas that are characterized by recurrent *PLAGL1* fusions and enriched for pediatric patients.

## Introduction

Ependymomas encompass a heterogeneous group of central nervous system (CNS) neoplasms that occur along the entire neuroaxis and can affect both children and adults ^1^. DNA methylation and gene expression profiling efforts in recent years have identified several molecular groups of ependymoma across different anatomic sites of the CNS with distinct clinicopathological characteristics and molecular alterations or patterns ^2-10^. Within the supratentorial compartment, two molecularly defined types of ependymoma are characterized by recurrent gene fusions, one involving the gene *ZFTA* (formerly referred to as *C11orf95*, most frequently fused to *RELA*), and the other involving *YAP1* ^2,5^. More recently, several reports have expanded on the spectrum of gene fusions observed in supratentorial ependymoma and ependymoma-like tumors, in particular in the pediatric setting ^11-13^. Implementing these molecular markers into the WHO classification for brain tumors is of paramount importance in overcoming the challenges of histologically diverse tumor types and in increasing diagnostic accuracy. Still, many cases do not fit into the as of yet established CNS tumor types, leaving clinicians and patients with unclear or even incorrect diagnoses in further decision making.

Genome-wide DNA methylation profiling has emerged as a powerful tool for both robust classification of known CNS tumor entities and identification of novel and clinically relevant subclasses of brain tumors with characteristic alterations ^2,14^. Here, we describe a molecularly distinct subset of supratentorial neoplasms (n = 40) with ependymal appearance identified by investigation of a large cohort of DNA methylation data. These tumors harbor recurrent fusions involving the *pleomorphic adenoma gene-like 1* (*PLAGL1*) gene.

## Materials and methods

### Sample collection

Tumor samples and retrospective clinical data from 40 patients were obtained from multiple national and international collaborating centers and collected at the Department of Neuropathology of the University Hospital Heidelberg (Germany). Sample selection was based on unsupervised visualization of genome-wide DNA methylation data that revealed a molecularly distinct group of tumors (n = 40) forming a cluster separate from all established entities. A proportion of data was generated in the context of the Molecular Neuropathology 2.0 study. Analysis of tissue and clinical data was performed in accordance with local ethics regulations. Clinical details of the patients are listed in Supplementary Table 1 (online resource).

### Histology and immunohistochemistry

For all cases with sufficient material (n = 16), histological review of an H&E-stained slide was performed according to the World Health Organization (WHO) 2016 classification of tumors of the CNS ^15^. Immunohistochemical staining was performed on a Ventana BenchMark ULTRA Immunostainer using the ultraView Universal DAB Detection Kit (Ventana Medical Systems, Tucson, AZ, USA). Antibodies were directed against: glial fibrillary acid protein (GFAP; Z0334, rabbit polyclonal, 1:1000 dilution, Dako Agilent, Santa Clara, CA, USA), epithelial membrane antigen (EMA; clone GP1.4, mouse monoclonal, dilution 1:1000, Thermo Fisher Scientific, Fremont, CA, USA), Sry-related HMG-BOX gene 10 (SOX10; clone EP268, rabbit monoclonal, dilution 1:100, Cell Marque Corp., Rocklin, CA, USA) and oligodendrocyte lineage transcription factor 2 (OLIG2; clone EPR2673, rabbit monoclonal, dilution 1:50, Abcam, Cambridge, UK).

### DNA methylation array processing and copy number profiling

Genome-wide DNA methylation profiling of all samples was performed using the Infinium MethylationEPIC (EPIC) BeadChip (Illumina, San Diego, CA, USA) or Infinium HumanMethylation450 (450k) BeadChip array (Illumina) according to the manufacturer’s instructions and as previously described ^14^. Raw data were generated at the Department of Neuropathology of the University Hospital Heidelberg, the Genomics and Proteomics Core Facility of the German Cancer Research Center (DKFZ) or at respective international collaborator institutes, using both fresh-frozen and formalin-fixed paraffin-embedded (FFPE) tissue samples. All computational analyses were performed in R version 3.6.0 (R Development Core Team, 2016; https://www.R-project.org). Copy-number variation analysis from 450k and EPIC methylation array data was performed using the conumee Bioconductor package version 1.12.0 ^16^. Raw signal intensities were obtained from IDAT-files using the minfi Bioconductor package version 1.21.4. Illumina EPIC and 450k samples were merged to a combined data set by selecting the intersection of probes present on both arrays (combineArrays function, minfi). Each sample was individually normalized by performing a background correction (shifting of the 5% percentile of negative control probe intensities to 0) and a dye-bias correction (scaling of the mean of normalization control probe intensities to 10,000) for both color channels. Subsequently, a correction for the array type (450k/EPIC) was performed by fitting univariable, linear models to the log2-transformed intensity values (removeBatchEffect function, limma package version 3.30.11). The methylated and unmethylated signals were corrected individually. Beta-values were calculated from the retransformed intensities using an offset of 100 (as recommended by Illumina). All samples were checked for duplicates by pairwise correlation of the genotyping probes on the 450k/EPIC array. To perform unsupervised non-linear dimension reduction, the remaining probes after standard filtering ^14^ were used to calculate the 1-variance weighted Pearson correlation between samples. The resulting distance matrix was used as input for t-SNE analysis (t-distributed stochastic neighbor embedding; Rtsne package version 0.13). The following non-default parameters were applied: theta = 0, pca = F, max_iter = 10,000 perplexity = 20.

### RNA sequencing and analysis

RNA was extracted from FFPE tissue samples using the automated Maxwell system with the Maxwell 16 LEV RNA FFPE Kit (Promega, Madison, WI, USA), according to the manufacturer’s instructions. Transcriptome analysis using messenger RNA (mRNA) sequencing of samples for which RNA of sufficient quality and quantity was available was performed on a NextSeq 500 instrument (Illumina) as previously described ^17^. This was possible for 20 tumors within the novel group and 14 *ZFTA:RELA*-fused ependymomas. In addition, a reference cohort of other glioma and glioneuronal subtypes were used for differential gene expression analysis (*YAP1:MAMLD1*-fused ependymoma (n = 3), central neurocytoma (n = 9), extraventricular neurocytoma (n = 8), dysembryoplastic neuroepithelial tumor (n = 11), papillary glioneuronal tumor (n = 9), *KIAA1549:BRAF*-fused pilocytic astrocytoma (n = 14), diffuse midline glioma H3 K27M mutant (n = 14) and glioblastoma IDH-wildtype (n = 9)). Fastq files from transcriptome sequencing were used for *de novo* annotation of fusion transcripts using the deFuse ^18^ and Arriba (v1.2.0) ^19^ algorithms with standard parameters. All further analysis was performed in R (version 3.6.0; R Core Team, 2019) using the DESeq2 package (v1.28.1) ^20^. Principal Component Analysis (PCA) was performed after variance stabilizing transformation of the count data and normalization with respect to library size, based on the selection of the top 1,000 most variable genes with relative log expression normalization. Similarities between samples were determined by computing Manhattan distances on the variance stabilized data followed by unsupervised hierarchical clustering. Differential expression testing was performed on raw count data after fitting a negative binomial model. P-values were adjusted for multiplicity by applying the Benjamini-Hochberg correction.

### Targeted next generation DNA sequencing and mutational analysis

Genomic DNA was extracted from FFPE tumor tissue samples of 18 patients within the cohort using the automated Maxwell system with the Maxwell 16 FFPE Plus LEV DNA Purification Kit (Promega, Madison, WI, USA), according to the manufacturer’s instructions. Capture-based next-generation DNA sequencing was performed on a NextSeq 500 instrument (Illumina) as previously described ^21^ using a custom brain tumor panel (Agilent Technologies, Santa Clara, CA, USA) covering the entire coding and selected intronic and promoter regions of 130 genes of particular relevance in CNS tumors (Supplementary Table 2, online resource).

### Statistical analysis

Statistical analysis was performed using GraphPad Prism 9 (GraphPad Software, La Jolla, CA, USA). Data on survival could be retrospectively retrieved for ten patients. Distribution of time to progression or recurrence (TTP) after surgery was estimated by the Kaplan-Meier method and compared between groups with the log-rank test. Patients lost to follow-up are censored at date of last contact in analysis of TTP. P-values below 0.05 were considered significant.

## Results

### DNA methylation profiling reveals a molecular distinct group of ependymoma

DNA methylation profiling has emerged as a powerful approach for robust classification of CNS neoplasms ^14^. Using a screening approach based on unsupervised visualization of a large cohort of genome-wide DNA methylation data, we identified a highly distinct group of tumors (n = 40) forming a cluster separate from all established entities of which a high proportion of tumors (19/32, 59%) were histopathologically diagnosed as ependymoma. A more focused t-SNE analysis of DNA methylation patterns of these samples alongside 453 other well-characterized ependymal neoplasms (reference samples included in the current version of the Heidelberg DNA methylation classifier with a calibrated score >0.9) confirmed the distinct nature of this novel group (Fig. 1 and Supplementary Figure 1, online resource). Analysis of copy number variations (CNVs) derived from DNA methylation array data revealed a relatively balanced profile in most of the cases, with structural aberrations on chromosome 22q (21/40, 52.5%) and 6q (19/40, 47.5%) most frequently observed (Supplementary Figure 2a, online resource). A chromothripsis-like pattern affecting chromosomes 6 and 13 was seen in one of the samples (Supplementary Figure 2b, online resource). In one case a homozygous deletion of *CDKN2A/B* was detected. An integrated plot of CNVs identified in all samples is given in Supplementary Figure 2c (online resource).

**Fig. 1.**
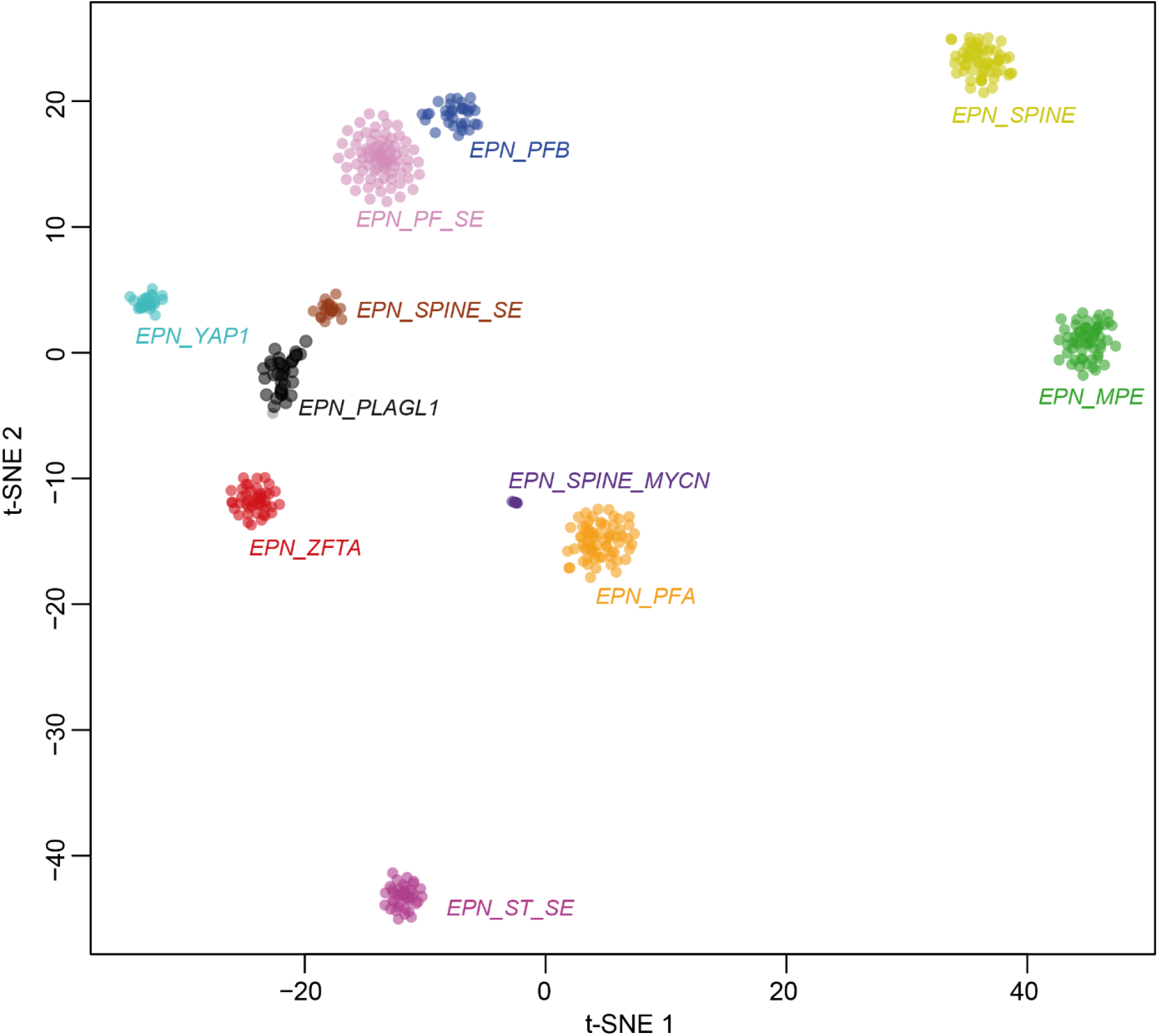
DNA methylation profiling reveals a molecular distinct group of ependymoma. t-distributed stochastic neighbor embedding (t-SNE) analysis of DNA methylation profiles of the 40 tumors investigated (*EPN_PLAGL1*) alongside selected reference samples. Reference DNA methylation classes: ependymoma posterior fossa group A (*EPN_PFA*), ependymoma posterior fossa group B (*EPN_PFB*), ependymoma spinal (*EPN_SPINE*), ependymoma with *ZFTA* fusion (*EPN_ZFTA*), ependymoma with *YAP1* fusion (*EPN_YAP1*), myxopapillary ependymoma (*EPN_MPE*), spinal ependymoma (*EPN_SPINE*), posterior fossa subependymoma (*EPN_PF_SE*), spinal subependymoma (*EPN_SPINE_SE*), supratentorial subependymoma (*EPN_ST_SE*) and spinal ependymoma with *MYCN* amplification (*EPN_SPINE_MYC*). Additional clustering analyses indicated that the PLAGL1 cohort can potentially be further subdivided into two clusters (not shown).

### Recurrent rearrangements involving *PLAGL1* are characteristic for the novel group of ependymoma

Since a high proportion of supratentorial ependymomas are driven by gene fusions involving *ZFTA* (*C11orf95*, most frequently fused to *RELA*) or *YAP1*, we performed mRNA sequencing of all samples with sufficient material (n = 20). In 19/20 of the cases, a gene fusion involving *PLAGL1* was detected, conserving the zinc finger structure (C2H2 type) as part of the fusion product, with either *EWSR1* as 5’ partner or *FOXO1* or *EP300* as a 3’ partner (Fig. 2a-c). In the most common *EWSR1:PLAGL1* fusions (n = 13), exons 1-9 or 1-8 of *EWSR1* (NM_013986), which is located on chromosome 22q12.2, were fused to exon 5 of *PLAGL1* (NM_001289039), which is found on chromosome 6q24.2. Five out of 20 cases were observed with exons 1-5 of *PLAGL1* fused to *FOXO1* upstream of exons 2-3 (NM_0017612) were also observed. In one case, exons 1-5 of *PLAGL1* are fused to exons 15-31 of *EP300* (NM_001429). In all rearrangements, the DNA binding domain (zinc finger structure) of *PLAGL1* was retained and fused to the respective transactivation domain (TAD) of the partner gene (Fig. 2a-c). We next performed an exploratory differential gene expression analysis of tumor samples (n = 20) within the novel group in comparison to *ZFTA:RELA*-fused ependymomas (n = 14). Unsupervised hierarchical clustering demonstrated a clear segregation of tumor samples in comparison to *ZFTA:RELA*-fused ependymoma (Fig. 2d). These results were recapitulated by PCA of normalized transcript counts (Fig. 2e). Quantification of mRNA expression revealed that the *PLAGL1* gene itself was more highly expressed in tumors within the novel group than in *ZFTA:RELA*-fused ependymoma (adjusted p = 1.22e-14; Fig. 2f and g). Additionally, upregulated genes of potential interest included *H19* and *IGF2* (adjusted p = 1.31e-83, adjusted p = 5.04e-08; Fig. 2h and i), both regulated by *PLAGL1* and with known functions in the tumorigenesis of different cancers ^22^. *RELA* and *ZFTA* transcript levels were upregulated in *ZFTA:RELA*-fused ependymomas (adjusted p = 1.03e-61 and adjusted p = 1.10e-19, respectively; Fig. 2j and k). Differential gene expression analysis between tumors within the novel group and a reference cohort of other glial and glioneuronal subtypes confirmed high transcript levels of *PLAGL1* (adjusted p = 2.35e-18), *H19* (adjusted p = 9.12e-15), *IGF2* (adjusted p = 7.91e-06) and *DLK1* (adjusted p = 1.12e-10) in the *PLAGL1*-fused cohort (Figure 3a-c and Supplementary Figure 3, online resource). Expression of particular markers differentially expressed in astrocytic and in ependymal neoplasms ^23-25^ revealed low *OLIG2* and *SOX10* expression (adjusted p = 3.89e-26 and adjusted p = 7.75e-65) within the novel group, with similar expression of *GFAP* (Figure 3d-f and Supplementary Figure 3, online resource). Analysis of the mutational landscape of 19/40 tumors in the novel group using targeted next-generation sequencing revealed *TERT* promoter mutations (C228T) in two of the cases (Supplementary Table 1, online resource), with no other relevant events involving putative brain tumor genes.

**Fig. 2.**
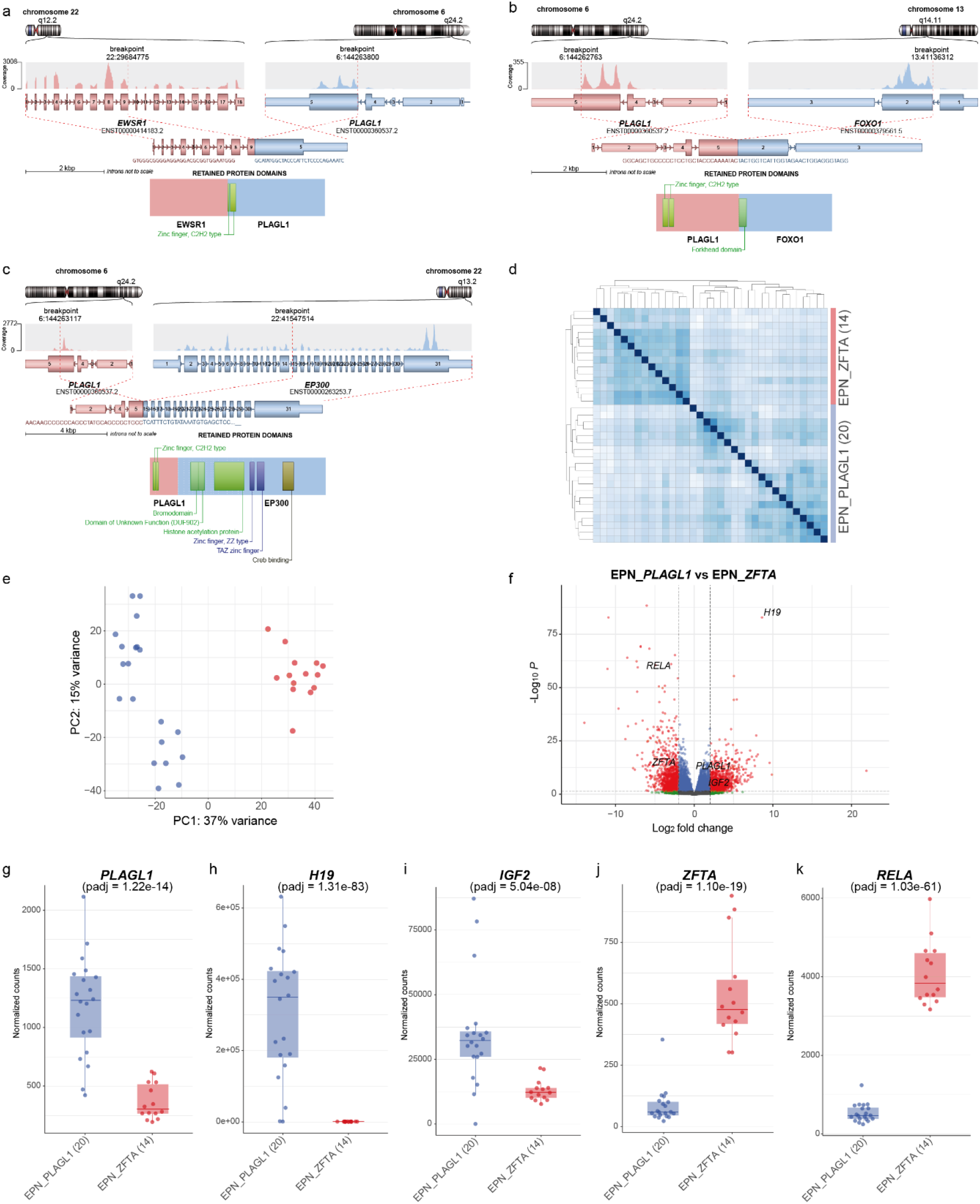
Illustration of the *PLAGL1* fusion genes and transcriptional profiling of tumors samples in the novel group. Visualization of the *PLAGL1* fusion genes detected by RNA sequencing for three selected samples. *EWSR1:PLAGL1* fusion in case #1, in which exons 1-9 of *EWSR1*, as the 5’ partner, are fused to exon 5 of *PLAGL1* (a), *PLAGL1:FOXO1* fusion in case #14, in which exons 1-5 of *PLAGL1* are fused to exons 2-3 of *FOXO1* as the 3’ partner (b), and *PLAGL1:EP300* fusion in case #19, in which exons 1-5 of *PLAGL1* are fused to exons 15-31 of *EP300* as the 3’ partner (c), conserving the zinc finger structure (C2H2 type) as part of the fusion products. Differences in gene expression profiles between samples in the novel group and *ZFTA:RELA*-fused ependymomas. Normalized transcript counts from samples in the novel group and *ZFTA:RELA*-fused ependymomas clustered by Pearson’s correlation coefficient (d) and principal component analysis (e). Volcano plot depicting genes differentially expressed between samples in the novel group versus *ZFTA:RELA*-fused ependymomas (f). *PLAGL1* (g), *H19* (h), *IGF2* (i), *ZFTA* (j), and *RELA* (k) expression in the novel group (n = 20) compared to *ZFTA:RELA*-fused ependymoma samples (n = 14).

**Fig. 3.**
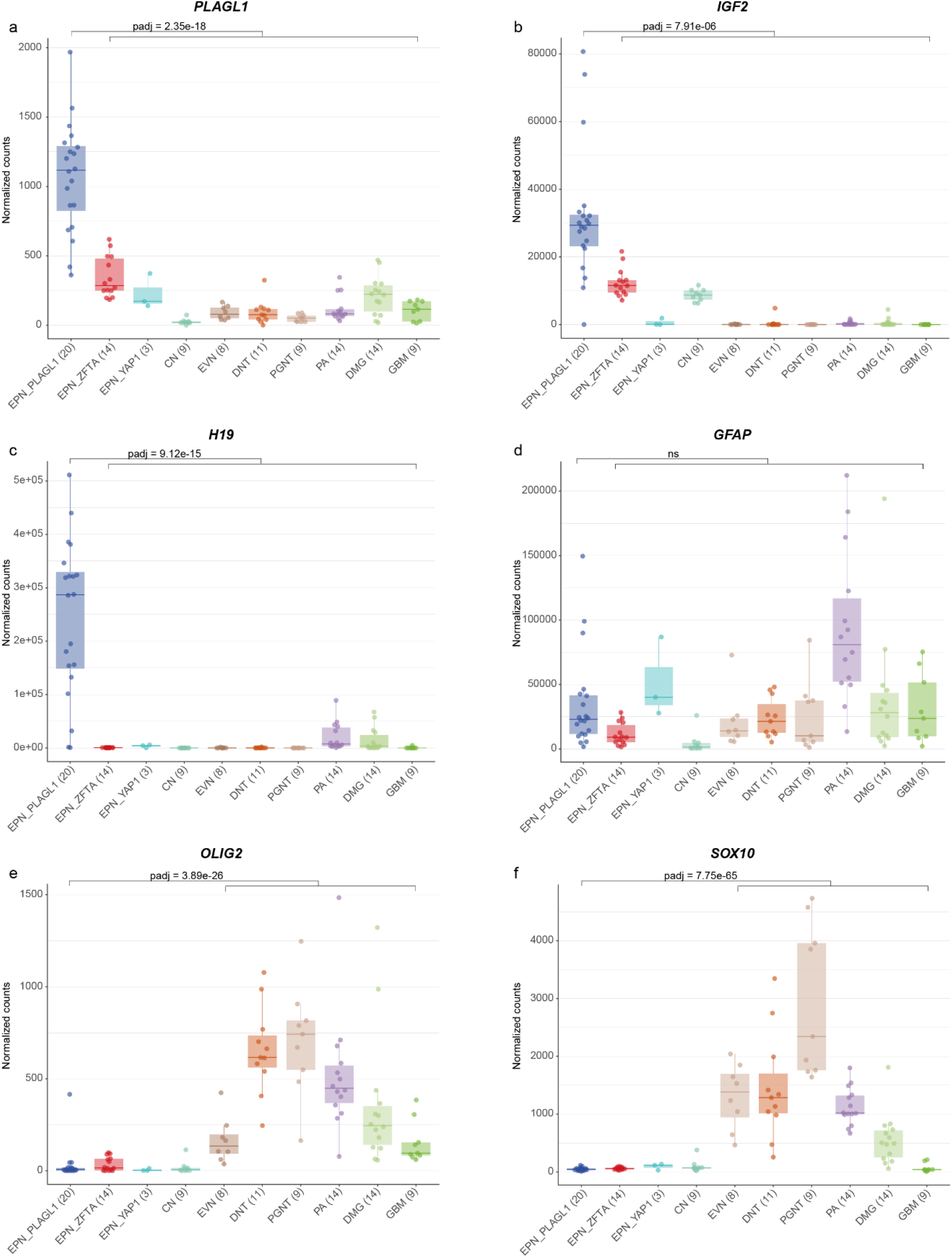
Transcriptional profiling of *PLAGL1*-altered ependymoma. Differential gene expression analysis between samples in the novel group (EPN_PLAGL1) and a reference cohort of different glial/glioneuronal tumors (*ZFTA:RELA*-fused ependymoma (EPN_ZFTA), *YAP1:MAMLD1*-fused ependymoma (EPN_YAP1), central neurocytoma (CN), extraventricular neurocytoma (EVN), dysembryoplastic neuroepithelial tumor (DNT), papillary glioneuronal tumor (PGNT), *KIAA1549:BRAF*-fused pilocytic astrocytoma (PA), diffuse midline glioma H3 K27M-mutant (DMG) and glioblastoma IDH-wildtype (GBM). *PLAGL1, IGF2* and *H19* are more highly expressed in EPN_PLAGL1 cases when compared with representative glial/glioneuronal tumors (a-c). *GFAP* levels are similar compared to different glial/glioneuronal tumors (d). Expression of markers differentially expressed in astrocytic and in ependymal tumors revealed low *OLIG2* and *SOX10* expression in EPN_PLAGL1 compared to astrocytic/glioneuronal tumors (e,f).

### Clinical characteristics and morphological features demonstrate pediatric-type tumors with ependymoma-like appearance

Analysis of available clinical data demonstrated that median age of the patients at the time of diagnosis was 6.2 years (n = 25; range 0-30; with 92% of the tumors occurring in patients < 17 years of age, Fig. 4a) and the sex distribution was relatively balanced (F/M = 1:1.2, Fig. 4b). All tumors in our series were located supratentorially (Fig. 4c). Outcome data were available for 11 patients. Analysis of time to recurrence in comparison to *ZFTA:RELA*-fused ependymomas (n = 80) revealed a similar outcome (p = 0.18; Figure 4d). The initial histopathological diagnoses of the tumors within the cohort were relatively wide, although a high proportion of cases were designated as ependymoma (19/32, 59%). Other recurrent diagnoses included ‘embryonal tumor’ and different low- and high-grade gliomas (Supplementary Table 1, online resource). More detailed descriptions of the cases are given in Supplementary Table 1. A histopathological review of samples with available material (n = 16) confirmed a relatively wide morphological spectrum of tumors with ependymoma-like features (Fig. 5a-i). Histologically, all reviewed tumors shared a moderate to high increase in cellular density in a mostly fine neurofibrillary matrix with prominent microcystic changes (Fig. 5a-d). The tumor cells typically had monomorphic, round to oval nuclei with finely dispersed chromatin and prominent nucleoli. Single cases presented more pleomorphic cells. In many cases, perivascular pseudorosettes were observed, at least focally. Two of the cases showed focal oligodendroglial morphology with perinuclear halos due to cytoplasmatic clearing (Fig. 5e). Extensive calcification was seen in a small number of tumors (n = 3). Necrosis was not observed. Mitotic activity was generally low, with exception of two cases. Immunoreactivity for GFAP was present in all cases (Fig. 5f). The tumor cells neither expressed OLIG2 nor SOX10 (Fig. 5g and h). In 3/16 of the cases, a dot-like positivity for EMA was detected (Fig. 5i).

**Fig. 4.**
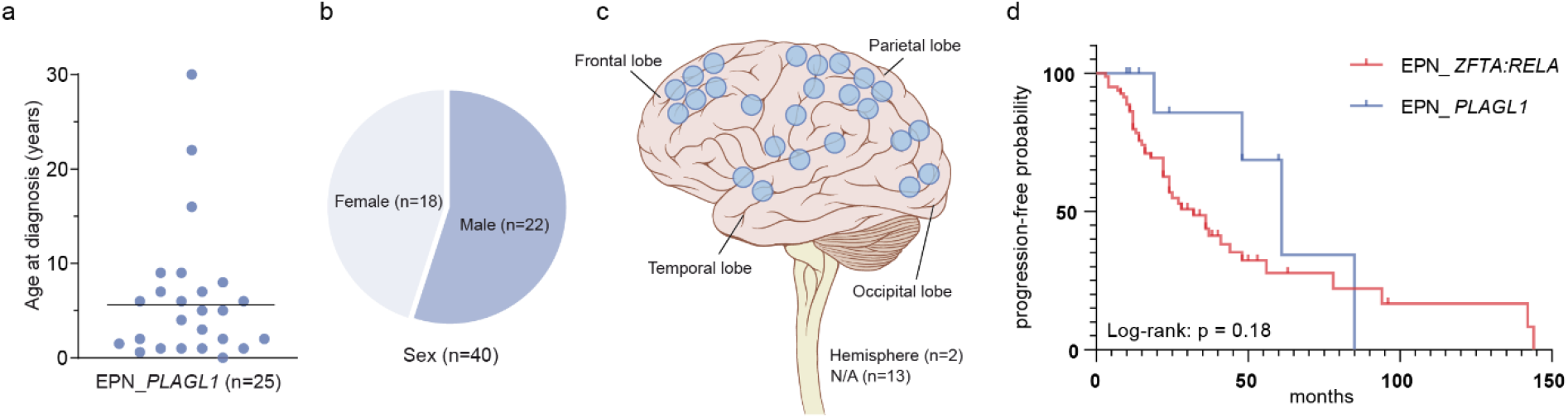
Clinical features of the investigated cohort. Age at diagnosis with the median age of 6.2 years (a), patient sex distribution (b) and distribution of tumor location (c). Time to progression or recurrence (TTP) of 11 patients from the investigated cohort (EPN_*PLAGL1*) for whom follow-up data were available compared to TTP of 80 patients with *ZFTA:RELA*-fused ependymoma (EPN_*ZFTA:RELA*; d).

**Fig. 5.**
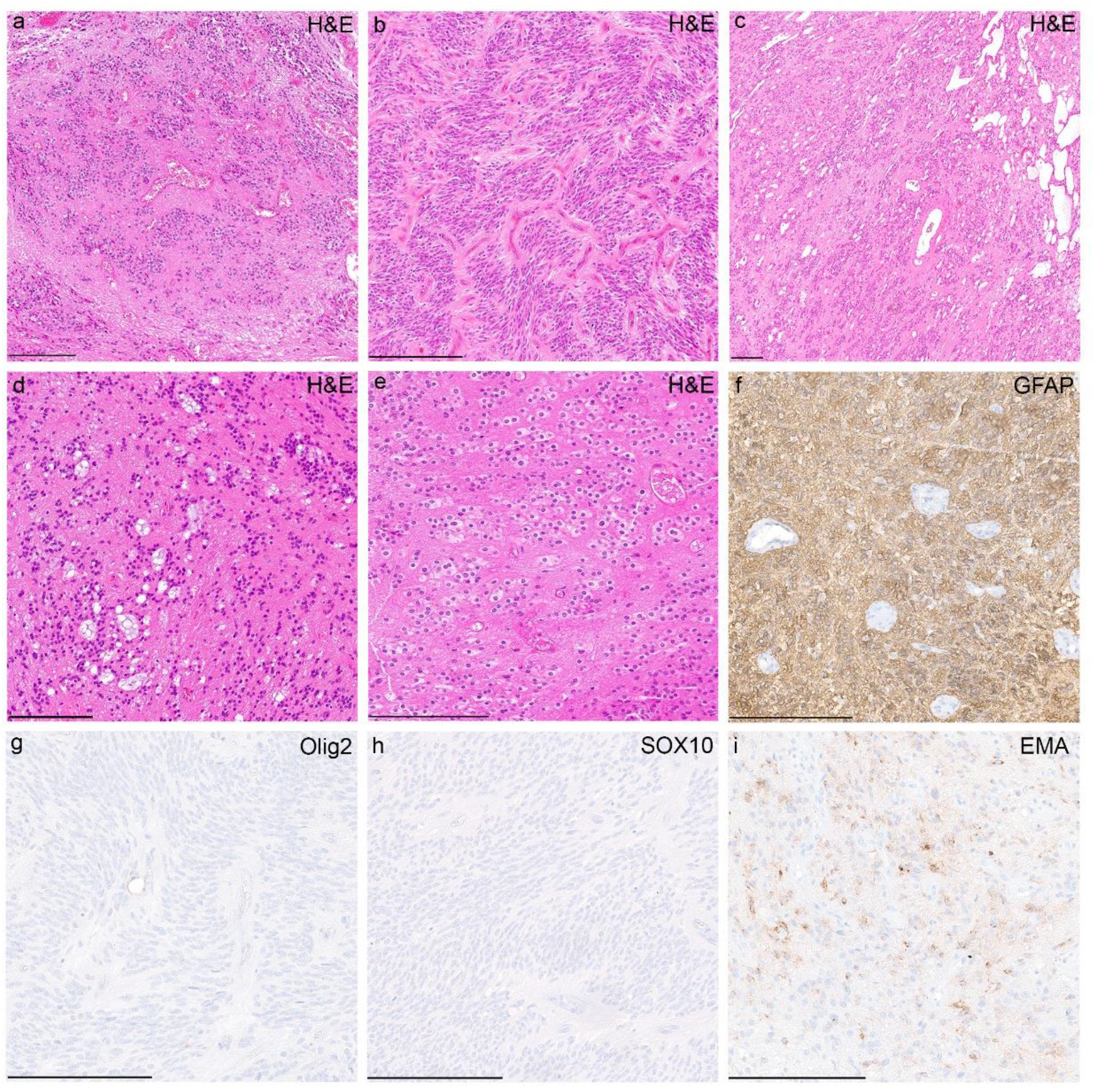
Morphological and immunohistochemical features of tumors within the cohort. Histologically, tumors shared a moderate to high increase in cellular density with mostly monomorphic, round to oval nuclei and often prominent microcystic changes (a-d). Perivascular pseudorosettes were observed in several of the cases, although very subtle in some the samples (a-d). Occasionally, tumor cells showed oligodendroglial morphology with perinuclear halos due to cytoplasmatic clearing (e). Immunohistochemically, tumors were GFAP-positive (f) and OLIG2- and SOX10-negative (g,h). In 3/16 of the cases, a dot-like positivity for EMA was detected (i). Scale bars 200 μm.

## Discussion

Here, we provide evidence for the pathobiological heterogeneity of ependymal tumors beyond the established spectrum by reporting the existence of an epigenetically distinct group of rare pediatric-type supratentorial neoplasms with often ependymoma-like appearance that shows recurrent gene fusions involving the *PLAGL1* gene.

Our findings suggest rearrangements involving *PLAGL1*, particularly *EWSR1:PLAGL1* and *PLAGL1:FOXO1* fusions, as a molecular hallmark of this novel group of tumors. Gene fusions of *PLAGL1* with *EWSR1* have been reported exceptionally rarely in neoplasms of the CNS, including single cases of a *SMARCB1*-deficient atypical teratoid/rhabdoid tumor (AT/RT) ^26^ and a glioneuronal tumor, not elsewhere classified (NEC) ^27^. However, in a very recent report, a *PLAGL1:EWSR1* fusion was described in a supratentorial ependymoma of a six-year-old child ^11^. While *EWSR1* has long been known to be involved in gene fusions in Ewing sarcoma and several other tumor entities ^28^, the role of *PLAGL1* in tumorigenesis is not yet fully understood. The *PLAGL1* gene encodes a C2H2 zinc finger protein that acts as a transcription factor as well as cofactor of other regulatory proteins, and is expressed in diverse types of human tissues amongst others in neural stem/progenitor cells and developing neuroepithelial cells ^29,30^. Although its specific role in tumorigenesis is controversial and its functions appear to depend on the cellular context, altered expression of *PLAGL1* has been linked to various types of cancer ^31-33^. More recent studies provide evidence for its oncogenic function in brain tumors with overexpression of *PLAGL1* being involved in tumorigenesis of glioblastoma ^34,35^ and interaction of *PLAGL* family transcription factors in *ZFTA:RELA*-fused supratentorial ependymoma ^36^.

In the *EWSR1:PLAGL1* fusions described here, the whole N-terminal transcriptional activation domain (TAD) of *EWSR1* is fused in-frame to the zinc finger domain (with DNA binding activity) of *PLAGL1*, very similar to other oncogenic *EWSR1* fusions, in particular rearrangements between *EWSR1* and *PATZ1* ^*37,38*^. This indicates aberrant recruitment of the TAD of *EWSR1* to the DNA binding domain of *PLAGL1* with subsequent downstream effects, as described for other *EWSR1* rearrangements, as the likely oncogenic function of this fusion ^39^. This also fits to the increased expression of *PLAGL1* in these samples. In addition, five samples harbored a fusion between *PLAGL1* and the transcriptional factor *FOXO1*, which is a known partner in other rearrangements ^40,41^. In the *PLAGL1:FOXO1* fusion observed here, the DNA binding domain of *PLAGL1* is juxtaposed to the C-terminal TAD of *FOXO1*, which seems quite similar to *PAX3:FOXO1* rearrangements as frequently observed in alveolar rhabdomyosarcoma ^41^. In a single case, *PLAGL1* was fused to *EP300*, a fusion partner known from ‘CNS tumors with *BCOR* alteration’ ^42^. Additionally, upregulated genes included *H19* and *IGF2*, both regulated by *PLAGL1* and with known functions in tumorigenesis of different cancers ^22^. This might indicate a potential downstream effect of the fusion. However, the precise oncogenic mechanism of the EWSR1:PLAGL1, PLAGL1:FOXO1 and PLAGL1:EP300 chimeric proteins remain to be elucidated. Further studies will be needed to reveal the exact role of the fusions in these tumors.

Another important finding was the relatively wide morphological spectrum of tumors within this group. Although most tumors were originally diagnosed as ependymoma, a significant proportion of cases were designated to other entities, including different low- and high-grade tumors. Consistent with that, a histopathological review of cases with sufficient material revealed a morphologically heterogeneous group of tumors often with ependymoma-like features, compatible with the designation as ependymoma. These findings are supported by differential gene expression analysis between tumors within the novel group and a reference cohort of other glial and glioneuronal tumors, that revealed low expression levels of *OLIG2* and *SOX10*, both suggested to distinguish astrocytic from ependymal tumors ^23-25^. However, the absence of a unifying morphological pattern in this group of tumors underlines the relevance of molecular profiling for precise diagnosis of these CNS neoplasms. This group has not been identified as a distinct subset in previous large-scale ependymoma studies due to the relatively small case numbers, broad morphology and lack of routine RNA profiling in previous cohorts, again highlighting the importance of RNA sequencing in standard brain tumor diagnostic. According to the structure of specifying ‘essential diagnostic criteria’ of the upcoming 5th edition of the WHO classification of CNS tumors, we suggest (a) the specific signature by DNA methylation profiling or (b) the combination GFAP expression and *PLAGL1* fusions as essential diagnostic criteria for these tumors.

A limitation of our study is the relatively low extent of clinical data, in particular patient outcome data, which allows only a rough estimation of the malignancy of the tumors within this novel group. Considering the high number of cases without sequencing data, it seems also possible that other alterations apart from the described fusions could be present, particularly in those tumors which do not show indication for a *PLAGL1* fusion in the copy-number profile. Follow-up analyses are needed to characterize this new group of CNS neoplasms in more detail.

In summary, we provide new insights into the tumorigenesis of ependymoma and identify *PLAGL1* as a putative relevant driver in this entity. Given their ependymoma-like appearance and expression pattern with a high frequency of *PLAGL1* fusions, we suggest the term ‘supratentorial ependymoma with *PLAGL1* fusion’ to describe this novel group of tumors. These findings have immediate implications for brain tumor profiling in order to avoid incorrect diagnoses due to lack of alignment with established tumor types. *PLAGL1*-fusion positive ependymomas should thus be included into upcoming classifications of brain tumors.

## Acknowledgements

For excellent technical support we sincerely thank the Microarray Unit of the German Cancer Research Center (DKFZ) Genomics and Proteomics Core Facility, as well as I. Leis and M. Schalles (Department of Neuropathology, Institute of Pathology, University Hospital Heidelberg, Heidelberg, Germany). We also thank the German Childhood Cancer Foundation for funding (“Neuropath 2.0 - Increasing diagnostic accuracy in paediatric neurooncology” (DKS 2015.01)). This study was supported by the Hertie Network of Excellence in Clinical Neuroscience. P. Sievers is a fellow of the Hertie Academy of Excellence in Clinical Neuroscience. S. C. Henneken received scholarships of the Mildred-Scheel doctoral program of the German Cancer Aid and the German Academic Scholarship Foundation. S. Brandner was partly funded by the National Institute of Health Research (NIHR) UCLH/UCL Biomedical Research Centre. A subset of the human tissue was obtained from University College London NHS Foundation Trust as part of the UK Brain Archive Information Network (BRAIN UK, Ref: 19/001) which is funded by the Medical Research Council and Brain Tumour Research UK. U. Schüller was supported by the Fördergemeinschaft Kinderkrebszentrum Hamburg.

## Author contributions

P.S., S.C.H., K.W.P. and F.S. designed the study; P.S., S.C.H., D.T.W.J., K.W.P. and F.S. conducted experiments; P.S., S.C.H., C.B., D.Sc., D.St., K.O., D.T.W.J., K.W.P. and F.S. analyzed data; all authors assisted with material and data collection; all authors contributed to manuscript writing.

## References

1 Louis DN, O. H., Wiestler OD, Cavenee WK. WHO classification of tumours of the central nervous system. Vol. Revised 4 edn. (IARC, Lyon, 2016).

2 Pajtler, K. W. et al. Molecular Classification of Ependymal Tumors across All CNS Compartments, Histopathological Grades, and Age Groups. Cancer Cell 27, 728–743, doi:10.1016/j.ccell.2015.04.002 (2015).

3 Pajtler, K. W. et al. Molecular heterogeneity and CXorf67 alterations in posterior fossa group A (PFA) ependymomas. Acta Neuropathol 136, 211–226, doi:10.1007/s00401-018-1877-0 (2018).

4 Panwalkar, P. et al. Immunohistochemical analysis of H3K27me3 demonstrates global reduction in group-A childhood posterior fossa ependymoma and is a powerful predictor of outcome. Acta Neuropathol 134, 705–714, doi:10.1007/s00401-017-1752-4 (2017).

5 Parker, M. et al. C11orf95-RELA fusions drive oncogenic NF-kappaB signalling in ependymoma. Nature 506, 451–455, doi:10.1038/nature13109 (2014).

6 Ghasemi, D. R. et al. MYCN amplification drives an aggressive form of spinal ependymoma. Acta Neuropathol 138, 1075–1089, doi:10.1007/s00401-019-02056-2 (2019).

7 Cavalli, F. M. G. et al. Heterogeneity within the PF-EPN-B ependymoma subgroup. Acta Neuropathol 136, 227–237, doi:10.1007/s00401-018-1888-x (2018).

8 Witt, H. et al. Delineation of two clinically and molecularly distinct subgroups of posterior fossa ependymoma. Cancer Cell 20, 143–157, doi:10.1016/j.ccr.2011.07.007 (2011).

9 Witt, H. et al. DNA methylation-based classification of ependymomas in adulthood: implications for diagnosis and treatment. Neuro Oncol 20, 1616–1624, doi:10.1093/neuonc/noy118 (2018).

10 Wani, K. et al. A prognostic gene expression signature in infratentorial ependymoma. Acta Neuropathol 123, 727–738, doi:10.1007/s00401-012-0941-4 (2012).

11 Zschernack, V. et al. Supratentorial ependymoma in childhood: more than just RELA or YAP. Acta Neuropathol, doi:10.1007/s00401-020-02260-5 (2021).

12 Pages, M. et al. Diagnostics of pediatric supratentorial RELA ependymomas: integration of information from histopathology, genetics, DNA methylation and imaging. Brain Pathol 29, 325–335, doi:10.1111/bpa.12664 (2019).

13 Tomomasa, R. et al. Ependymoma-like tumor with mesenchymal differentiation harboring C11orf95-NCOA1/2 or -RELA fusion: A hitherto unclassified tumor related to ependymoma. Brain Pathol, e12943, doi:10.1111/bpa.12943 (2021).

14 Capper, D. et al. DNA methylation-based classification of central nervous system tumours. Nature 555, 469–474, doi:10.1038/nature26000 (2018).

15 Louis, D. N. et al. The 2016 World Health Organization Classification of Tumors of the Central Nervous System: a summary. Acta Neuropathol 131, 803–820, doi:10.1007/s00401-016-1545-1 (2016).

16 Aryee, M. J. et al. Minfi: a flexible and comprehensive Bioconductor package for the analysis of Infinium DNA methylation microarrays. Bioinformatics 30, 1363–1369, doi:10.1093/bioinformatics/btu049 (2014).

17 Stichel, D. et al. Routine RNA sequencing of formalin-fixed paraffin-embedded specimens in neuropathology diagnostics identifies diagnostically and therapeutically relevant gene fusions. Acta Neuropathol 138, 827–835, doi:10.1007/s00401-019-02039-3 (2019).

18 McPherson, A. et al. deFuse: an algorithm for gene fusion discovery in tumor RNA-Seq data. PLoS Comput Biol 7, e1001138, doi:10.1371/journal.pcbi.1001138 (2011).

19 Uhrig, S. et al. Accurate and efficient detection of gene fusions from RNA sequencing data. Genome Res 31, 448–460, doi:10.1101/gr.257246.119 (2021).

20 Love, M. I., Huber, W. & Anders, S. Moderated estimation of fold change and dispersion for RNA-seq data with DESeq2. Genome Biol 15, 550, doi:10.1186/s13059-014-0550-8 (2014).

21 Sahm, F. et al. Next-generation sequencing in routine brain tumor diagnostics enables an integrated diagnosis and identifies actionable targets. Acta Neuropathol 131, 903–910, doi:10.1007/s00401-015-1519-8 (2016).

22 Varrault, A. et al. Zac1 regulates an imprinted gene network critically involved in the control of embryonic growth. Dev Cell 11, 711–722, doi:10.1016/j.devcel.2006.09.003 (2006).

23 Kleinschmidt-DeMasters, B. K. et al. SOX10 Distinguishes Pilocytic and Pilomyxoid Astrocytomas From Ependymomas but Shows No Differences in Expression Level in Ependymomas From Infants Versus Older Children or Among Molecular Subgroups. J Neuropathol Exp Neurol 75, 295–298, doi:10.1093/jnen/nlw010 (2016).

24 Ligon, K. L. et al. The oligodendroglial lineage marker OLIG2 is universally expressed in diffuse gliomas. J Neuropathol Exp Neurol 63, 499–509, doi:10.1093/jnen/63.5.499 (2004).

25 Otero, J. J., Rowitch, D. & Vandenberg, S. OLIG2 is differentially expressed in pediatric astrocytic and in ependymal neoplasms. J Neurooncol 104, 423–438, doi:10.1007/s11060-010-0509-x (2011).

26 Ramkissoon, S. H. et al. Clinical targeted exome-based sequencing in combination with genome-wide copy number profiling: precision medicine analysis of 203 pediatric brain tumors. Neuro Oncol 19, 986–996, doi:10.1093/neuonc/now294 (2017).

27 Lopez-Nunez, O. et al. The spectrum of rare central nervous system (CNS) tumors with EWSR1-non-ETS fusions: experience from three pediatric institutions with review of the literature. Brain Pathol 31, 70–83, doi:10.1111/bpa.12900 (2021).

28 Thway, K. & Fisher, C. Mesenchymal Tumors with EWSR1 Gene Rearrangements. Surg Pathol Clin 12, 165–190, doi:10.1016/j.path.2018.10.007 (2019).

29 Valente, T., Junyent, F. & Auladell, C. Zac1 is expressed in progenitor/stem cells of the neuroectoderm and mesoderm during embryogenesis: differential phenotype of the Zac1-expressing cells during development. Dev Dyn 233, 667–679, doi:10.1002/dvdy.20373 (2005).

30 Vega-Benedetti, A. F. et al. PLAGL1: an important player in diverse pathological processes. J Appl Genet 58, 71–78, doi:10.1007/s13353-016-0355-4 (2017).

31 Abdollahi, A. LOT1 (ZAC1/PLAGL1) and its family members: mechanisms and functions. J Cell Physiol 210, 16–25, doi:10.1002/jcp.20835 (2007).

32 Godlewski, J. et al. PLAGL1 (ZAC1/LOT1) Expression in Clear Cell Renal Cell Carcinoma: Correlations with Disease Progression and Unfavorable Prognosis. Anticancer Res 36, 617–624 (2016).

33 Su, H. C. et al. Gene expression profiling identifies the role of Zac1 in cervical cancer metastasis. Sci Rep 10, 11837, doi:10.1038/s41598-020-68835-0 (2020).

34 Hide, T. et al. Sox11 prevents tumorigenesis of glioma-initiating cells by inducing neuronal differentiation. Cancer Res 69, 7953–7959, doi:10.1158/0008-5472.CAN-09-2006 (2009).

35 Li, C. et al. Tumor edge-to-core transition promotes malignancy in primary-to-recurrent glioblastoma progression in a PLAGL1/CD109-mediated mechanism. Neurooncol Adv 2, vdaa163, doi:10.1093/noajnl/vdaa163 (2020).

36 Arabzade, A. et al. ZFTA-RELA Dictates Oncogenic Transcriptional Programs to Drive Aggressive Supratentorial Ependymoma. Cancer Discov, doi:10.1158/2159-8290.CD-20-1066 (2021).

37 Siegfried, A. et al. EWSR1-PATZ1 gene fusion may define a new glioneuronal tumor entity. Brain Pathol 29, 53–62, doi:10.1111/bpa.12619 (2019).

38 Rossi, S. et al. Expanding the spectrum of EWSR1-PATZ1 rearranged CNS tumors: An infantile case with leptomeningeal dissemination. Brain Pathol, e12934, doi:10.1111/bpa.12934 (2020).

39 Krystel-Whittemore, M. et al. Novel and established EWSR1 gene fusions and associations identified by next-generation sequencing and fluorescence in-situ hybridization. Hum Pathol 93, 65–73, doi:10.1016/j.humpath.2019.08.006 (2019).

40 Antonescu, C. R. et al. Novel GATA6-FOXO1 fusions in a subset of epithelioid hemangioma. Mod Pathol, doi:10.1038/s41379-020-00723-4 (2020).

41 Linardic, C. M. PAX3-FOXO1 fusion gene in rhabdomyosarcoma. Cancer Lett 270, 10–18, doi:10.1016/j.canlet.2008.03.035 (2008).

42 Tauziede-Espariat, A. et al. The EP300:BCOR fusion extends the genetic alteration spectrum defining the new tumoral entity of “CNS tumors with BCOR internal tandem duplication”. Acta Neuropathol Commun 8, 178, doi:10.1186/s40478-020-01064-8 (2020).

