## Supplementary material for "Recurrent fusions in *PLAGL1* define a distinct subset of pediatric-type supratentorial ependymoma": Suppl. Figures

**Supplementary Fig. 1** DNA methylation profiling reveals a molecular distinct group of ependymoma. t-distributed stochastic neighbor embedding (t-SNE) analysis of DNA methylation profiles of the 40 tumors investigated (*EPN_PLAGL1*) alongside selected reference samples. Reference DNA methylation classes: ependymoma posterior fossa group A (*EPN_PFA*), ependymoma posterior fossa group B (*EPN_PFB*), ependymoma spinal (*EPN_SPINE*), ependymoma with *ZFTA* fusion (*EPN_ZFTA*), ependymoma with *YAP1* fusion (*EPN_YAP1*), myxopapillary ependymoma (*EPN_MPE*), spinal ependymoma (*EPN_SPINE*), posterior fossa subependymoma (*EPN_PF_SE*), spinal subependymoma (*EPN_SPINE_SE*), supratentorial subependymoma (*EPN_ST_SE*) and spinal ependymoma with *MYCN* amplification (*EPN_SPINE_MYC*), pleomorphic xanthoastrocytoma (*PXA*), posterior fossa pilocytic astrocytoma (*PA_PF*), midline pilocytic astrocytoma (*PA_MID*), pilocytic astrocytoma and ganglioglioma (*PA/GG*), ganglioglioma (*GG*), rosette-forming glioneuronal tumor (*RGNT*), dysembryoplastic neuroepithelial tumor (*DNT*), extraventricular neurocytoma (*EVN*), papillary glioneuronal tumor (*PGNT*), diffuse leptomeningeal glioneuronal tumor subclass 1 and 2 (*DLGNT_1/2*), glioblastoma IDH-wildtype subclass mesenchymal (*GBM_MES*), glioblastoma IDH-wildtype subclass RTK I (*GBM_RTK I*), glioblastoma IDH-wildtype subclass RTK II (*GBM_RTK II*), glioblastoma IDH-wildtype H3.3 G34 mutant (*GBM_G34*) and diffuse midline glioma H3 K27M mutant (*DMG_K27*).

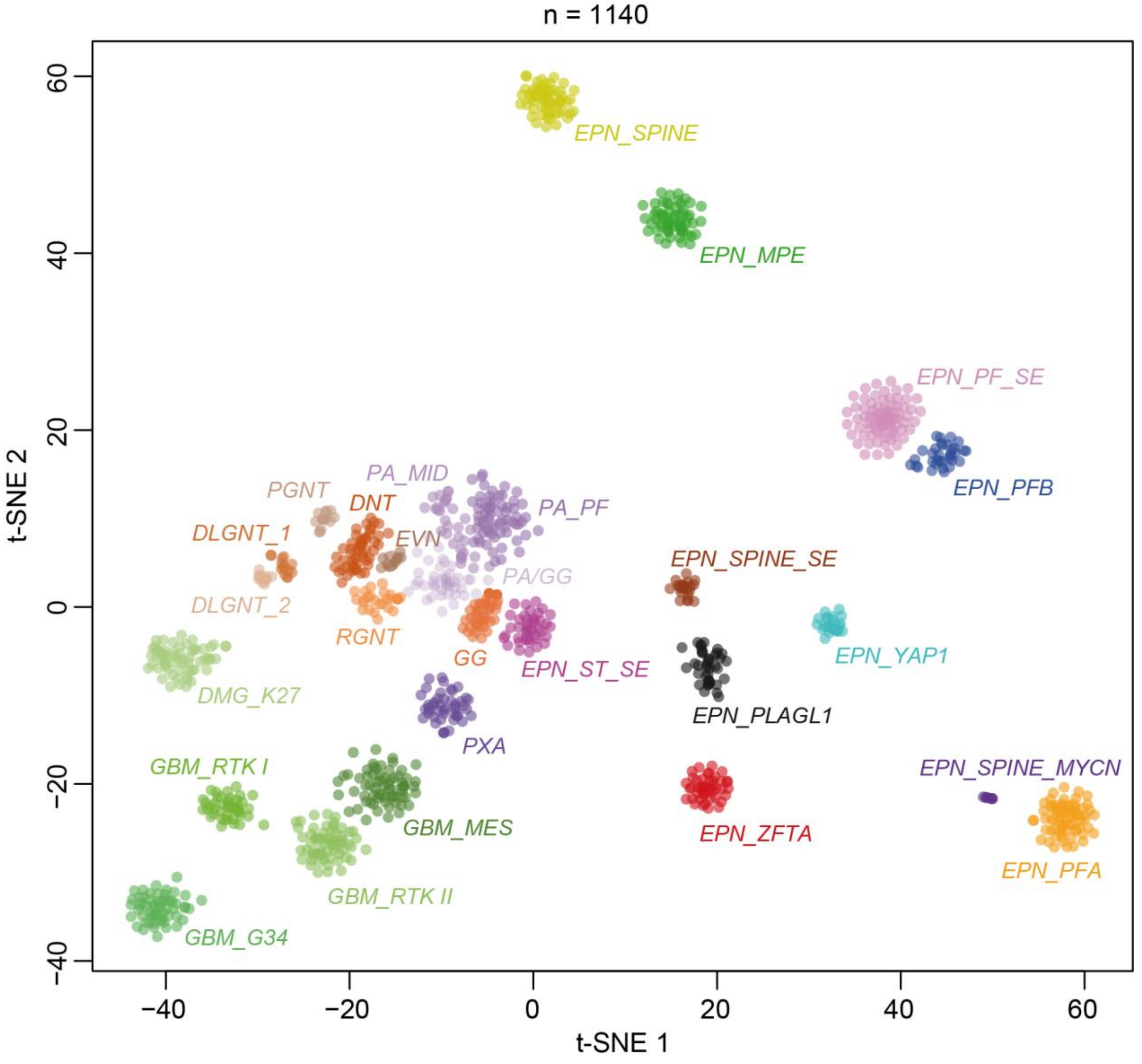

**Supplementary Fig. 2** Copy-number profiles in *PLAGL1*-fused ependymoma. Copy-number profiles derived from DNA methylation array of different tumors within the novel group showing structural alterations affecting chromosome 6q around the *PLAGL1* locus and chromosome 22q including *EWSR1* (a) as well as a chromothripsis-like pattern affecting chromosomes 6 and 13 (b). Integrated plot of copy number variations in all samples within the cohort show no recurrent chromosomal alterations besides small structural aberrations on chromosome 22q and 6q (c). The probes of the array are combined in 8000 bins (green/red dots). Gains/amplifications represent positive, losses represent negative deviations from the baseline.

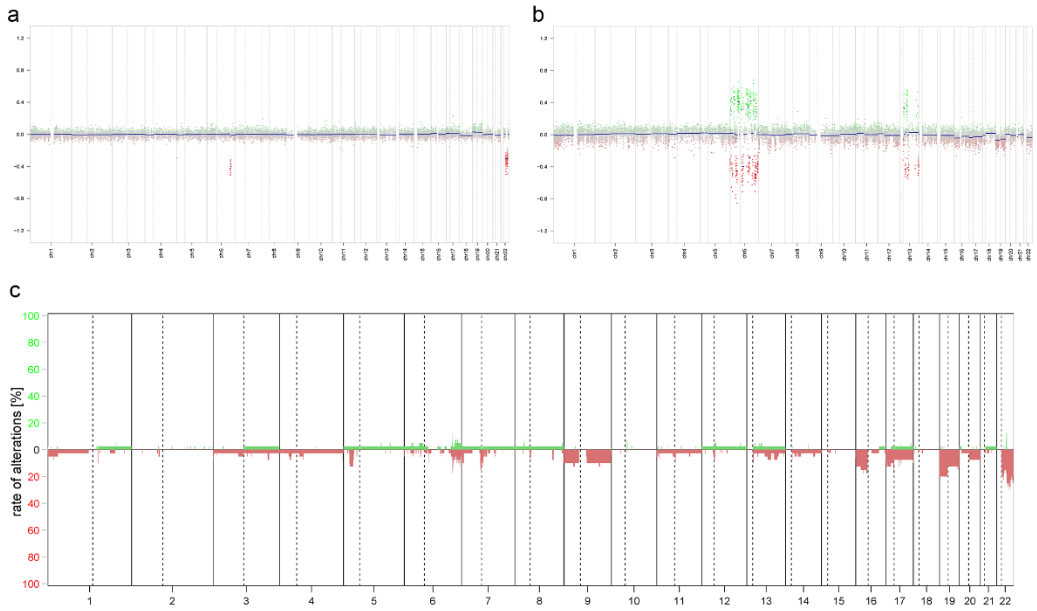

**Supplementary Fig. 3** Differences in gene expression profile between *PLAGL1*-fused ependymomas and different glial/glioneuronal tumors. Volcano plot depicting genes differentially expressed between samples in the novel group (EPN_PLAGL1) versus pilocytic astrocytoma (PA; a), dysembryoplastic neuroepithelial tumor (DNT; b), and glioblastoma IDH-wildtype (GBM; 3). *PLAGL1* and the imprinted genes *IGF2*, *H19* and *DLK1* are more highly expressed in EPN_PLAGL1 cases when compared with representative glial/glioneuronal tumors (a-c). Low *OLIG2* and *SOX10* expression levels in EPN_PLAGL1 compared to glial/glioneuronal tumors (a-c).

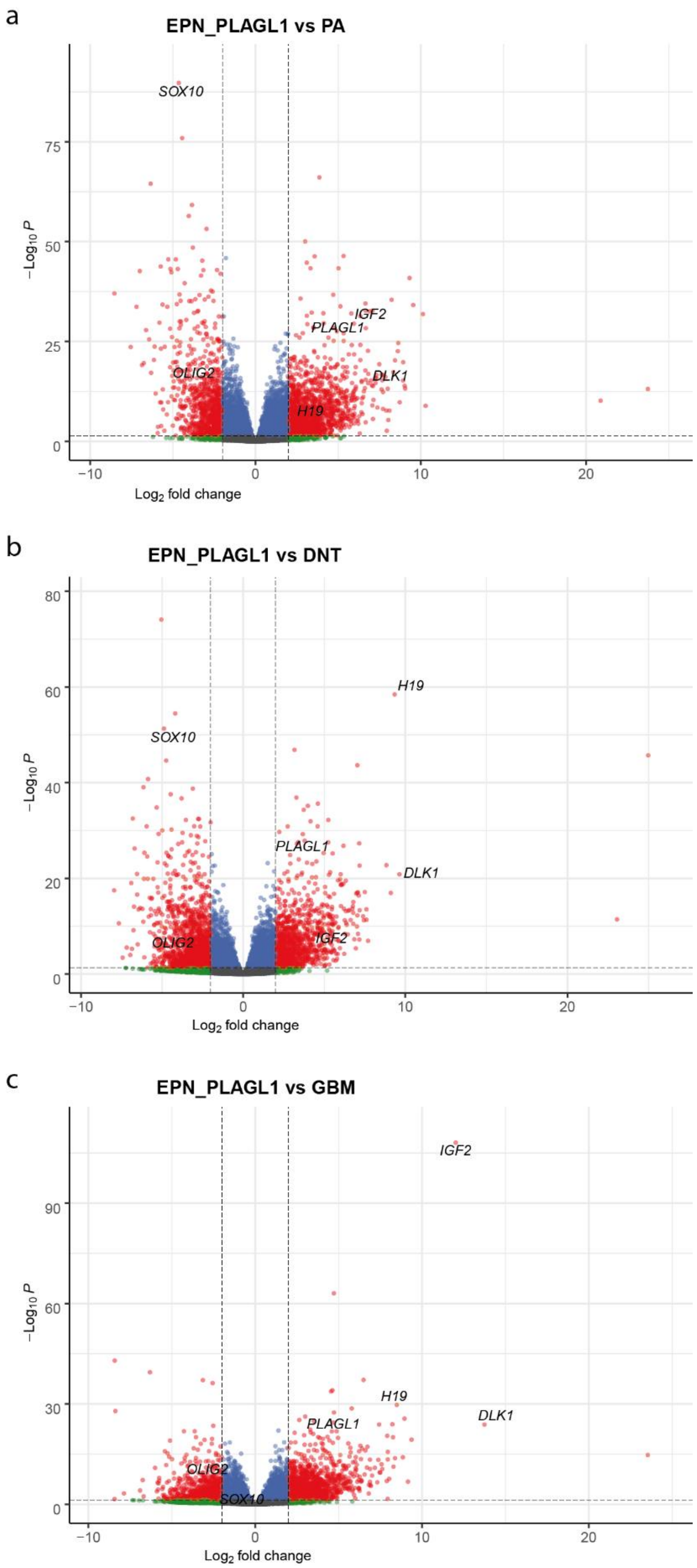
